# mBACS enhances pseudouridine profiling through efficient and robust chemical conversion

**DOI:** 10.64898/2026.09.05.749584

**Authors:** Feng Feng, Giuseppina Pisignano, Linzhen Kong, Thomas Grimes, Paul E. Brennan, Chun-Xiao Song

## Abstract

Pseudouridine (Ψ) is the most abundant RNA modification and regulates RNA stability, splicing, and translation. Utilizing the dual nucleophilic N^1^ and O^2^ of Ψ, we previously developed 2-bromoacrylamide-assisted cyclization sequencing (BACS) for quantitative, single-base Ψ detection. Here, we screened new dual electrophiles and developed methyl 2-bromoacrylate-assisted cyclization sequencing (mBACS), which achieves higher Ψ conversion efficiency and lower false-positive rates, enabling more sensitive and robust Ψ detection. The enhanced sensitivity of mBACS uncovered new Ψ sites in human tRNAs. mBACS further revealed TRUB1 and PUS10 as the exclusive redundant writers of the conserved tRNA Ψ55 modification and uncovered crosstalk between Ψ55 and other tRNA modifications. It also revealed 5-fluorouracil as a pan-pseudouridine synthase inhibitor that induces widespread but site-specific pseudouridylation remodelling. Finally, mBACS supported robust Ψ profiling from as little as 10 ng of total RNA, establishing it as a sensitive, quantitative, and low-input second-generation platform.

## Introduction

Ψ, often referred to as the “fifth nucleotide”, was the first RNA modification to be identified and remains the most abundant RNA modification in cellular RNAs. It is particularly enriched in constitutively expressed non-coding RNAs (ncRNAs), including ribosomal RNA (rRNA), small nuclear RNA (snRNA), and transfer RNA (tRNA). Ψ is installed by pseudouridine synthases (PUSs), a family of enzymes comprising 13 members annotated in the human genome^1^. Compared with uridine, Ψ contains an additional N-H group that enables the formation of an extra hydrogen bond, thereby enhancing RNA structural stability and modulating interactions with RNA-binding proteins^2^.

RNA pseudouridylation regulates a wide range of biological processes^3,4^. In rRNA, Ψ contributes to ribosome biogenesis and translational fidelity, whereas in tRNA it stabilizes RNA structure and promotes accurate codon recognition^5^. At U2 RNA, Ψ facilitates the assembly of early splicing complexes by strengthening interactions between U2 and the RNA helicase Prp5, which in turn enhances its ATPase activity and promotes efficient pre-mRNA splicing^3^. The functional significance of Ψ also extends to cancer, where altered Ψ patterns have been associated with different cancer types^4^, while aberrant expression of PUS enzymes often correlates with increased tumour aggressiveness and resistance to therapy^6^. Beyond its endogenous biological roles, Ψ has become a key component of mRNA vaccines, where it modulates the immune response, reduces innate immune sensing, enhances mRNA stability and translational efficiency, improving delivery and reducing side effects^7,8^. These diverse biological functions underscore the need for accurate and quantitative methods for mapping pseudouridylation.

To date, several chemical approaches have been developed for Ψ profiling. Among these, CMC-based methods - such as Pseudo-seq^9^, Ψ-seq^10^, PSI-seq^11^, and CeU-seq^12^ – rely on the selective reaction of Ψ with N-cyclohexyl-N′-(2-morpholinoethyl) carbodiimide methyl-p-toluenesulfonate (CMC)^9–12^, which introduces reverse transcription stops at modified sites. Bisulfite-based sequencing strategies, including RBS-Seq, BID-seq, and PRAISE^13–15^, introduce deletion signals at modified sites. However, CMC-based methods are semi-quantitative and bisulfite-based methods struggle to detect Ψ in densely modified regions and consecutive uridine sequences. Similarly, direct RNA sequencing using Oxford Nanopore technology offers a promising alternative for Ψ detection without chemical treatment^16,17^; however, modification calling remains computationally challenging and is currently limited by variable accuracy and sequence context-dependent signal interpretation.

To address these limitations, we recently developed BACS, which enables quantitative and accurate mapping of Ψ modification at the single-base resolution in cellular samples^18^. BACS exploited the dual nucleophilic N^1^ and O^2^ of Ψ and employed 2-bromoacrylamide as a dual electrophile, which first reacts with the free N^1^ of Ψ through Michael addition, followed by intramolecular alkylation of O^2^ of Ψ (Figure 1a, top). The resulting cyclized product enables the direct Ψ to C conversion, overcoming the limitations of RT-stop- and deletion-based methods and allowing accurate quantification of Ψ in densely modified regions and consecutive uridine sequences. Using BACS, we reported the first comprehensive substrate map of human PUS enzymes in tRNA^19^. Despite its broad utility, BACS still suffers from incomplete conversion, elevated false-positive rates in certain sequence motifs, and variable performance, likely due in part to the hygroscopic nature of 2-bromoacrylamide, which complicates storage and reduces reagent consistency over time.

**Figure 1:**
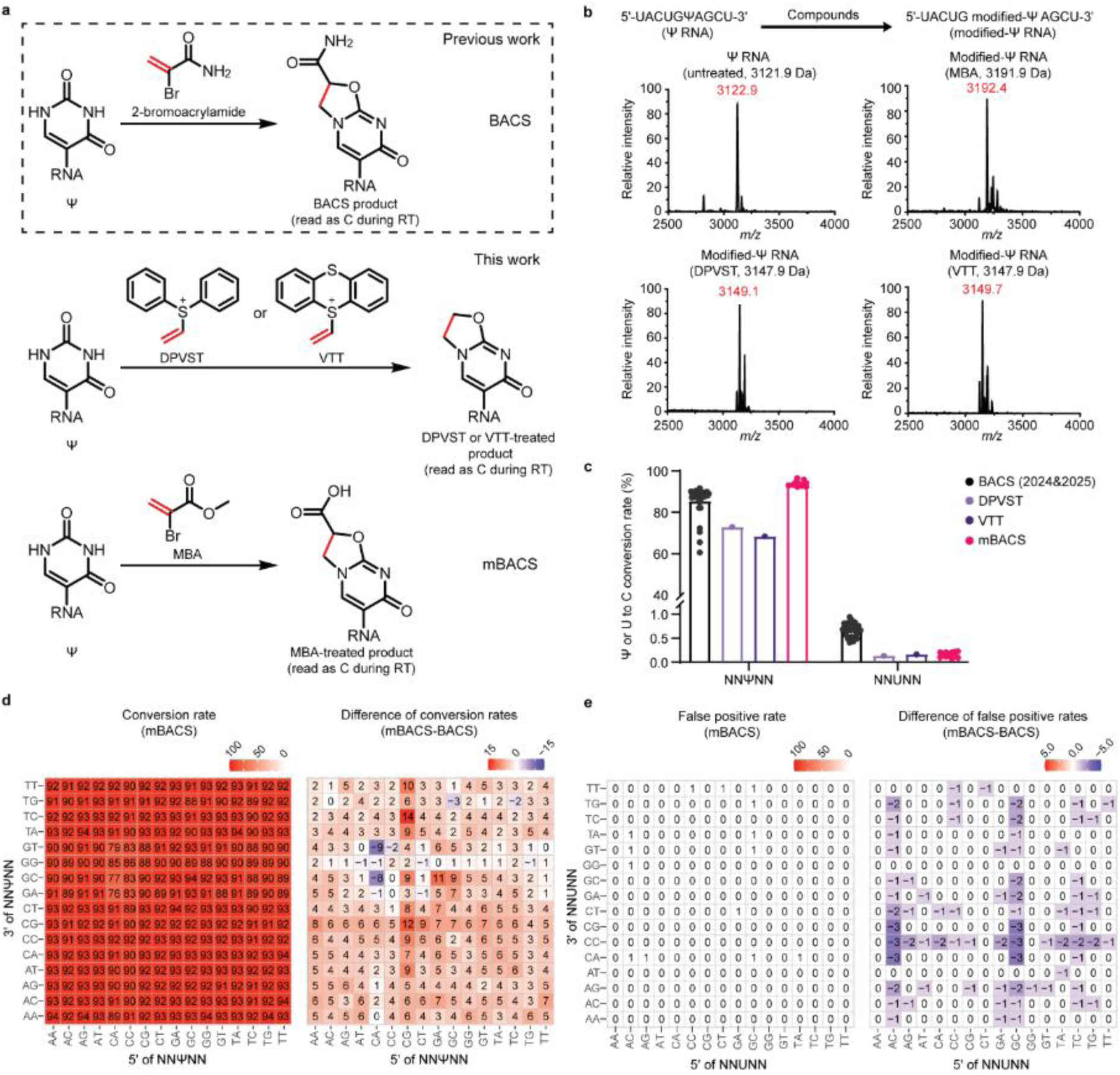
Chemical features and performance comparison of DPVST, VTT, and MBA for Ψ detection. a, Schematic illustration of the reactions between a Ψ-containing RNA oligonucleotide and the original BACS reagent (top), DPVST (middle), or VTT (middle), and MBA (bottom). b, MALDI-TOF mass spectra of the reaction products generated from MBA (top, right), DPVST (bottom, left), or VTT (bottom, right) following reaction with a 10-mer Ψ-containing RNA oligonucleotide. Calculated masses indicated in black, and detected masses indicated in red. c, Ψ/U to C conversion rates of the 30-mer NNΨNN and NNUNN spike-in RNAs following treatment with the original BACS reagent, MBA (mBACS), DPVST, and VTT. d, Conversion rates of the 30-mer NNΨNN spike-in RNA obtained using mBACS (left) and the difference in conversion rates with the previously reported BACS method (right). e, False positive rates of the 30-mer NNUNN spike-in RNA in mBACS (left) and the difference in false positive rates with those obtained using the previously reported BACS method (right). BACS data reported in c-e refer to previous dataset^18^.

To improve the sensitivity and robustness of Ψ detection, we evaluated alternative dual electrophiles including novel sulfonium compounds and other α-halo Michael acceptors. Sequencing analyses demonstrated that methyl 2-bromoacrylate (MBA) achieved higher conversion efficiency, lower false positive rates, and more consistent performance than BACS, enabling mBACS to faithfully recapitulate previously established Ψ landscapes in rRNAs and tRNAs while improving detection sensitivity and identifying additional Ψ sites with low modification levels. This enabled the identification of previously undetectable Ψ sites in both mitochondrial (mt) and cytoplasmic (cy) tRNAs, including Ψ65 in mt-tRNA and Ψ44 in cy-tRNA, likely installed by PUS1 and PUS7L, respectively. Application of mBACS to CRISPR-Cas9-mediated double-knockout models demonstrated that the conserved tRNA Ψ55 modification is redundantly installed by TRUB1 and PUS10, establishing these enzymes as the exclusive writers of this modification. Furthermore, mBACS uncovered crosstalk between Ψ55 and four additional cytoplasmic tRNA modifications, 3-amino-3-carboxypropyluridine at position 20 (acp³U20), N², N²- dimethylguanosine at position 26 (m²_2_G26), 3-methylcytidine at positions 20 and 32 (m³C20/32), and 1- methyladenosine at position 58 (m¹A58), revealing coordinated regulation of tRNA modification biogenesis. mBACS further characterized the pseudouridylation remodeling upon 5-fluorouracil (5-FU) treatment and revealed 5-FU as a pan-PUS inhibitor. Finally, mBACS demonstrated its suitability for robust Ψ profiling from as little as 10 ng of total RNA, expanding its applicability to low-input samples. Together, these advances establish mBACS as a sensitive, robust, and broadly applicable second- generation method for quantitative transcriptome-wide Ψ profiling.

## Results and Discussion

### Sulfonium compounds and MBA enable efficient and quantitative Ψ detection, with MBA showing superior performance

To address the limitation of the BACS method, we explored new dual electrophiles to improve Ψ- modification detection. Recently, sulfonium compounds such as diphenylvinylsulfonium triflate (DPVST) and vinylthianthrenium tetrafluoroborate (VTT) have emerged as efficient electrophilic probes for selective bioconjugation of amino acid residues under mild aqueous condition at room temperature^20,21^. Notably, their electrophilic properties enable chemical reactions with two distinct nucleophiles (Figure 1a, middle), resembling the chemical mechanism of BACS. We also tested other α-halo Michael acceptors including methyl 2-bromoacrylate (MBA) (Figure 1a, bottom) and 2-bromoacrylic acid (BAA).

We found that DPVST, VTT, and MBA efficiently reacted with a 10-mer Ψ-containing oligonucleotide. The corresponding cyclization products were successfully detected by matrix-assisted laser desorption/ionization mass spectrometry (MALDI) mass spectrometry, indicating that all three reagents have a chemical mechanism similar to 2-bromoacrylamide (Figure 1b). Unexpectedly, we observed an MBA-derived product of 14 Da mass loss. Subsequent NMR analysis of the reaction between MBA and a Ψ nucleoside revealed that the methyl ester group of MBA undergoes hydrolysis under basic conditions (pH 8.5) and elevated temperature (85°C) (Supplementary note 1). To further confirm the ester hydrolysis, we tested 10-mer Ψ-containing oligonucleotide with ethyl α-bromoacrylate (EBA) and observed a similar ester hydrolysis during the reaction process (Supplementary Figure 1a). Interestingly, BAA failed to react with Ψ (Supplementary Figure 1b), suggesting that the ester hydrolysis occurred after the initial Michael reaction and esterification of BAA is required for efficient reaction with Ψ.

We then compared the sequencing performance of DPVST, VTT, and MBA on synthetic 30-mer NNΨNN and NNUNN RNA spike-ins. Sequencing analysis showed that while DPVST and VTT exhibited lower conversion rates of 71% and 67% respectively than BACS (83.8%) at NNΨNN motifs, MBA achieved a higher conversion rate (92%) than BACS (Figure 1c and Supplementary Figure 1c-d). Importantly, all three compounds, MBA (0.14%), DPVST (0.12%), and VTT (0.15%), achieved lower false positive rates than BACS (0.64%) at NNUNN motifs (Figure 1c). Taken together, MBA achieved the highest conversion rate (Figure 1d) while maintaining a low false positive rate (Figure 1e), outperformed the other candidates and consistently provided a more robust performance than BACS (Figure 1c). Based on its superior performance, MBA was selected for all subsequent method validation and biological applications.

### mBACS faithfully recapitulates the BACS-derived Ψ landscape while improving detection sensitivity on low-stoichiometry Ψ sites

To validate the performance of mBACS, we applied the optimized mBACS workflow to total RNA isolated from HCT116 cells. Using a 5% modification-level threshold, we identified 108 high-confidence Ψ sites in human rRNA (Figure 2a). Notably, 106 of the 108 Ψ sites were also detected in the HCT116 dataset using the original BACS method^18^, whereas the remaining two Ψ sites with low modification levels, located at position 898 in 18S rRNA and 1795 in 28S rRNA (Figure 2b and Supplementary Figure 2a), were uniquely identified by mBACS, likely reflecting its improved conversion efficiency and reduced false-positive rate. Ψ4118 in 28S rRNA was not detected by mBACS because of its low modification level (Supplementary Figure 2a). Moreover, quantitative analysis showed a strong correlation in Ψ modification levels between the BACS and mBACS datasets, further confirming the robustness and reproducibility of the optimized mBACS method (Figure 2c).

**Figure 2:**
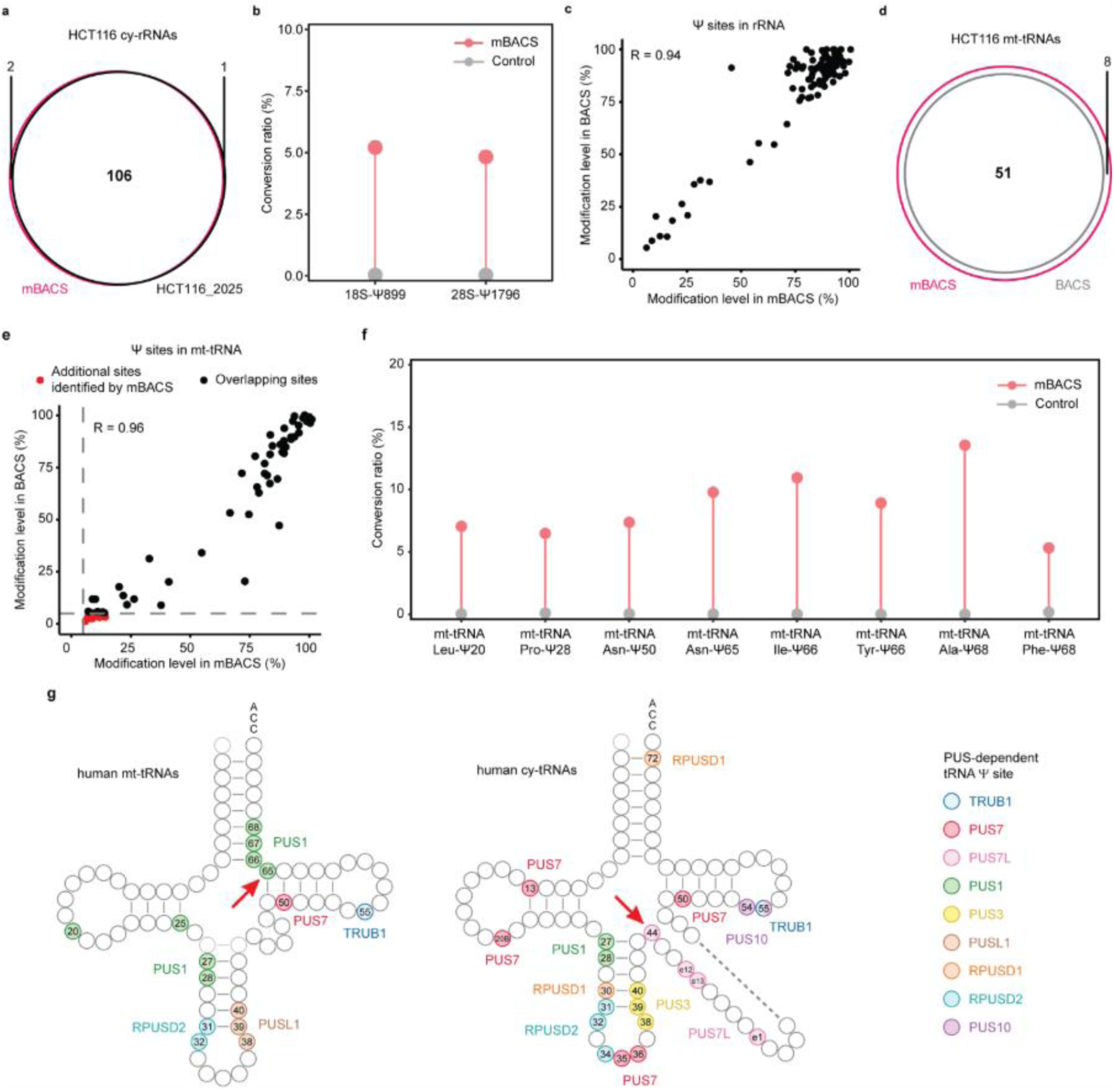
mBACS accurately recapitulates and expands rRNA and tRNA Ψ landscapes. a, Venn diagram showing the overlap of Ψ sites on cy-rRNAs detected by mBACS and the original BACS method in HCT116 cells. b, Conversion rates of Ψ898 and Ψ1795 sites on rRNA, mBACS treated group (red) and control group (grey). c, Scatter plot showing the correlation of Ψ modification levels at all identified sites in HCT116 cy-rRNAs between the BACS and mBACS methods. d, Venn diagram showing the overlap of Ψ sites identified in HCT116 mt-tRNAs by mBACS and the previously reported BACS method. e, Scatter plot showing the correlation of Ψ modification levels at all identified sites in HCT116 mt-tRNAs between the BACS and mBACS methods. f, Conversion rates of eight mBACS unique Ψ sites on mt-tRNAs in mBACS treated group (red) and control group (grey), including Leu-Ψ20, Pro-Ψ28, Asn-Ψ50, Asn-Ψ65, Ile-Ψ66, Tyr-Ψ66, Ala-Ψ68, and Phe-Ψ68. g, Comprehensive map of PUS- dependent Ψ sites in human mt-tRNAs (left) and cy-tRNAs (right). Red arrows indicate the newly identified sites mt-tRNA Ψ65 and cy-tRNA Ψ44. BACS data refer to previous dataset^18,19^.

Having validated mBACS for rRNA profiling, we next evaluated its performance on tRNAs, where our previous work established pseudouridylation as a major RNA modification^18^. We identified a total of 59 high-confidence Ψ sites in HCT116 mitochondrial tRNAs (mt-tRNAs) (Figure 2d), which showed strongest concordance with the previous BACS dataset (Figure 2e). Importantly, mBACS successfully detected all 51 high confidence Ψ sites previously identified by BACS in HCT116^19^. In addition, mBACS identified eight additional Ψ sites that were not classified as high-confidence in the previous BACS study because their modification levels fell below the 5% threshold (Figure 2f and Supplementary Figure 2b), primarily owing to lower conversion efficiency and/or elevated false-positive rates associated with specific sequence motifs, thereby demonstrating the improved sensitivity of mBACS.

To validate the eight additional Ψ sites identified by mBACS, we compared them with previously published Ψ datasets. Seven of them corresponded to known Ψ positions in mt-tRNAs from HeLa and HCT116 cells^18,19^, including Ψ68 in mt-tRNA-Ala and mt-tRNA-Phe, Ψ50 in mt-tRNA-Asn, Ψ66 in mt-tRNA-Ile and mt-tRNA-Tyr, Ψ20 in mt-tRNA-Leu (UUR), and Ψ28 in mt-tRNA-Pro (Figure 2f and Supplementary Figure 2b). The remaining site, Ψ65 in mt-tRNA-Asn, represents a previously unrecognized Ψ position in HCT116 cells. mBACS measured a modification level of 10.5%, compared with only 3.6% using BACS (Figure 2f and Supplementary Figure 2c). Consistent with this finding, Ψ65 was absent in PUS1-knockout HeLa cells in our previous BACS dataset^19^ (Supplementary Figure 2c), supporting PUS1 as the writer responsible for this modification.

We next applied mBACS to profile cytoplasmic tRNAs (cy-tRNAs) from HCT116 cells. Consistent with previous reports and our original BACS dataset^18,22^, mBACS detected the canonical Ψ positions in cy-tRNAs (Supplementary table 3). In addition, mBACS identified Ψ44, a previously unreported Ψ position that was neither detected by the original BACS workflow nor reported in previous studies. Its complete loss in PUS7L-knockout HCT116 cells in our previous BACS study further suggests PUS7L as the enzyme responsible for this modification^19^ (Supplementary Figure 2d).

Collectively, these results demonstrate that mBACS faithfully recapitulates known Ψ landscapes, while its enhanced sensitivity enables the detection of low-stoichiometry Ψ sites, including Ψ65 in mt-tRNAs and Ψ44 in cy-tRNAs, thereby expanding the detectable pseudouridylation landscape (Figure 2g) and establishing mBACS as a robust second-generation chemical sequencing platform for Ψ profiling.

### TRUB1/PUS10 double knockout establishes TRUB1 and PUS10 as the exclusive redundant Ψ55 writers and reveals functional crosstalk among cytoplasmic tRNA modifications

Our previous work identified TRUB1 and PUS10 as the primary PUS enzymes responsible for Ψ formation at position 55 in mt-tRNAs and position 54 in cy-tRNAs, respectively, while deletion of either TRUB1 or PUS10 alone resulted in only a modest reduction of Ψ55 in cy-tRNAs^19^. These observations suggested that TRUB1 and PUS10 function redundantly to catalyze Ψ55 formation in cy-tRNAs. However, it remained unknown whether TRUB1 and PUS10 are the exclusive PUS enzymes responsible for cy-tRNA Ψ55 installation and whether complete loss of this highly conserved modification could be achieved in human cells.

To address these questions, we generated TRUB1/PUS10 double-knockout (DKO) HCT116 using CRISPR-Cas9 and successfully isolated monoclonal cell lines lacking both TRUB1 and PUS10 enzymes (Supplementary Figure 3). Having established that mBACS accurately quantify Ψ54/55 in cy-tRNAs (Figure 3a), we next profiled pseudouridylation in DKO cells. Our results demonstrate that DKO completely abolishes Ψ55 formation in human cy-tRNAs, confirming that TRUB1 and PUS10 are the exclusive redundant PUS enzymes responsible for the conserved Ψ55 installation in human cy-tRNAs (Figure 3b).

**Figure 3:**
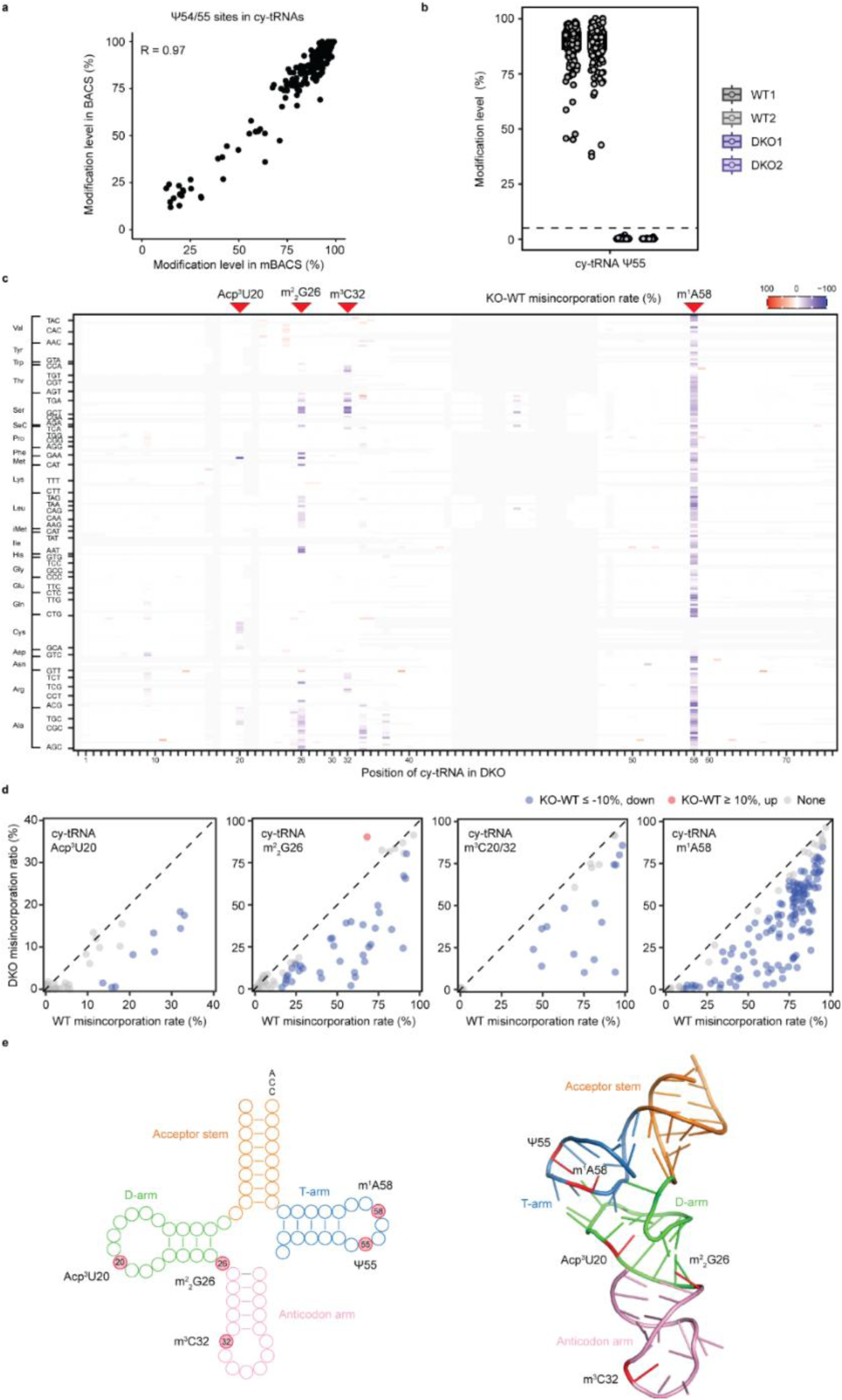
Complete loss of Ψ55 reveals TRUB1/PUS10 redundancy and functional crosstalk with additional cytoplasmic tRNA modifications. a, Scatter plot illustrating the correlation of modification levels at all identified Ψ54/55 sites in human cy- tRNAs between the BACS and mBACS methods. b, Ψ55 modification levels in cy-tRNAs between two HCT116 wild-type (WT) replicates and two DKO clones. Box plots show all Ψ sites at selected position; boxes represent the percentiles with a line at the median (Ψ55, n = 176 Ψ sites). c. Heatmap showing the differential misincorporation ratio of tRNA bases upon DKO. The x axis represents the canonical position of each aligned cy-tRNAs isodecoders. The y axis represents different cy-tRNA isodecoders. d. Scatter plot illustrating the misincorporation rate in acp³U20, m²_2_G26, m³C20/32 and m¹A58 in HCT116 WT and DKO. e, Cloverleaf secondary structure (left) and three-dimensional structure (right) of a tRNA, with each domain shown in a different color (Acceptor stem, orange; T arm, blue; D arm: green; Anticodon arm: pink), illustrating representative crosstalk between Ψ55 and four additional cytoplasmic tRNA modifications.

We next investigated whether complete loss of Ψ55 influences the installation of other tRNA modifications. We and others have previously shown that Ψ55 is installed at a very early stage of tRNA biogenesis, prior to tRNA 3’ end processing^19,23^. Several modifications that disrupt Watson-Crick base pairing, such as m¹A, m²_2_G, m³C, and acp³U, can be detected by characteristic reverse transcription misincorporation signatures during sequencing^24^. Neither TRUB1 KO nor PUS10 KO substantially altered the misincorporation rates of these modifications in cy-tRNAs (Supplementary Figure 4a-d). In contrast, the DKO markedly reduces the misincorporation signal corresponding to m^1^A58, a modification located in the same T-loop of tRNAs as Ψ55 (Figure 3c-e). Interestingly, this dependence of m^1^A58 on Ψ55 in cy-tRNA has also been reported in *S. cerevisiae*^23^, suggesting that the interplay between these two modifications is evolutionarily conserved. In addition to m^1^A58, the misincorporation signals corresponding to several other canonical tRNA modifications, including acp^3^U20, m^2^_2_G26, and m^3^C20/32, were significantly reduced in DKO compared to wild-type cells (Figure 3c-e), indicating that the efficient installation of these modifications depends on Ψ55. Notably, the misincorporation signals for these modifications were markedly reduced rather than completely abolished, suggesting that Ψ55 promotes proper tRNA folding or facilitates recognition by the corresponding modification enzymes.

Together, these findings establish TRUB1 and PUS10 as the exclusive redundant PUS enzymes responsible for Ψ55 installation in human cy-tRNA. Moreover, it reveals that Ψ55 is a key determinant for efficient installation of multiple other tRNA modifications, uncovering an evolutionarily conserved layer of coordination during tRNA maturation.

### mBACS reveals 5-FU as a pan-PUS inhibitor

5-FU has been a widely used chemotherapeutic agent for the treatment of multiple solid tumors for nearly 70 years. In addition to its canonical inhibition of thymidylate synthase, it has been shown to disrupt ribosome biogenesis and translation by impairing pre-rRNA processing and altering rRNA modification^25–28^. More recently, 5-FU has been shown to inhibit some PUS enzymes, such as PUS1, and reduce global RNA pseudouridylation, particularly in tRNAs, although these studies lacked single-base resolution and transcriptome-wide information^29,30^. We therefore investigated whether mBACS could resolve site-specific pseudouridylation remodelling following 5-FU treatment.

We treated HCT116 cells with 10 μM 5-FU for 24, 48, or 72 h and monitored cell growth throughout the treatment period. Cell growth was progressively inhibited, with the greatest effect observed after 72 h (Supplementary Figure 5a). Cells treated for 72h were then subjected to quantitative Ψ profiling using mBACS.Our results show that 5-FU treatment reduces Ψ modification levels in 28S, 5.8S, and 18S cy- rRNA, with a median decrease of approximately 10% across all detected Ψ sites in HCT116 (Figure 4a). Further analysis revealed substantial site-specific difference in sensitivity to 5-FU (Figure 4b-c). The inhibitory effects of 5-FU ranged from remarkable reduction (>30%) at some sites to little or no change at others (Figure 4b-c). These results indicate that although 5-FU broadly suppresses pseudouridylation in cytoplasmic rRNAs, individual Ψ sites differ markedly in their susceptibility to inhibition.

**Figure 4:**
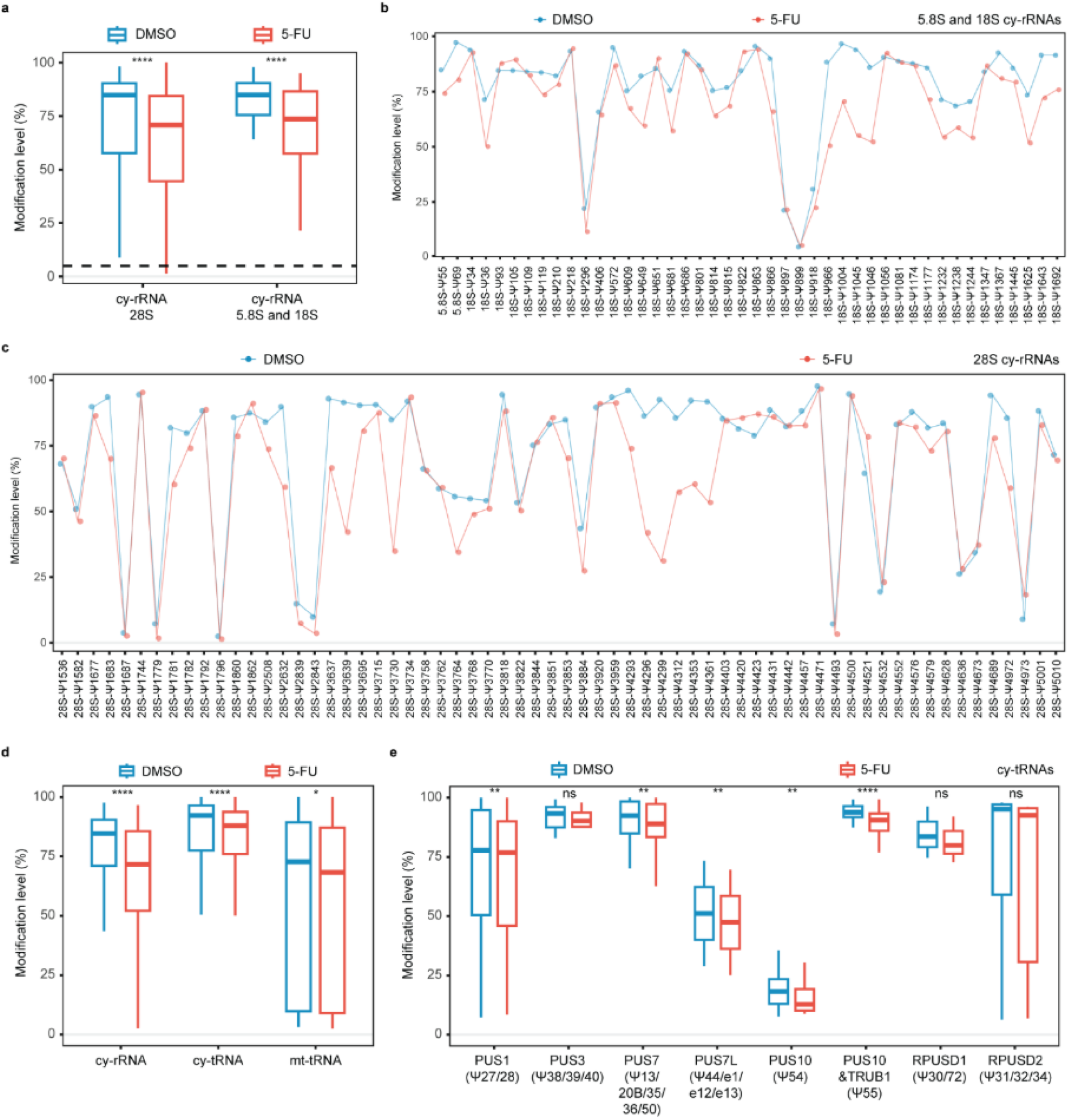
5-FU treatment induces selective remodeling of pseudouridylation in HCT116 rRNAs and tRNAs. a, Modification levels at all identified sites in HCT116 28S or 5.8S and 18S cy-rRNAs between DMSO and 5-FU treated group. Boxes represent the percentiles with a line at the median (28S rRNA: n = 128 Ψ sites; 5.8S and 18S: n=88 Ψ sites). b, Modification levels of each Ψ sites in HCT116 5.8S and 18S cy- rRNAs between DMSO and 5-FU treated group. c, Modification levels of each Ψ sites in HCT116 28S cy-rRNAs between DMSO and 5-FU treated group. d, Modification levels at identified sites in HCT116 cy-rRNA, cy-tRNAs, and mt-tRNA between DMSO and 5-FU treated group. Boxes represent the percentiles with a line at the median (cy-rRNA: n=108 Ψ sites; cy-tRNA: n=167 Ψ sites; mt-tRNA: n=57 Ψ sites). e, Modification levels at identified sites in HCT116 cy-tRNAs between DMSO and 5-FU treated group. Boxes represent the percentiles with a line at the median (PUS1: n=25 Ψ sites; PUS3: n=16 Ψ sites; PUS7: n=36 Ψ sites; PUS7L: n=2; PUS10: n=13 Ψ sites; Ψ55: n=63 Ψ sites; RPUSD1: n=3 Ψ sites; RPUSD2: n=9 Ψ sites). *P* values were calculated using paired, two-tailed t-test. ns, *P* >= 0.05; * *P* < 0.05; ** *P* < 0.01; *** *P* < 0.001 and **** *P* < 0.0001.

In addition to its effect on cy-rRNA, 5-FU treatment also caused an overall reduction of Ψ level on HCT116 cy-tRNAs (Figure 4d). The most pronounced inhibition was observed at Ψ55 (Figure 4e), catalyzed redundantly by TRUB1 and PUS10, as well as at the PUS10-dependent Ψ54 site. These results suggest that stand-alone PUS enzymes are also susceptible to 5-FU-mediated inhibition. Notably, the extent of pseudouridylation loss differed among RNA classes, with cy-rRNAs exhibiting the greatest reduction, followed by cy-tRNAs, whereas mt-tRNAs and mt-rRNAs were comparatively less affected by 5-FU treatment (Figure 4d and Supplementary Figure 5b-c).

Taken together, these results suggest that 5-FU acts as a pan-PUS inhibitor that induces widespread but highly site-specific remodeling of RNA pseudouridylation, revealing marked differences in the susceptibility of individual Ψ sites, RNA classes, and PUS enzymes to inhibition.

### mBACS enabled robust Ψ profiling from as little as 10 ng of total RNA

The standard mBACS workflow requires 100 ng of total RNA for rapid, accurate, and quantitative Ψ profiling. To further extend the applicability of mBACS to RNA-limited biological and clinical samples, we next investigated whether the method could be adapted for low-input applications. We reduced the RNA input tenfold, from 100 ng to 10 ng total RNA (approximately equivalent to RNA recovered from ∼500 HCT116 cells), and assessed the performance of mBACS. Despite the tenfold reduction in input material, mBACS maintained robust sequencing performance (Figure 5a). Specifically, the spike-in controls in the 10 ng mBACS group achieved a conversion rate of 92.4%, while maintaining a low false- positive rate (0.16%) comparable to that observed under standard input conditions (Figure 5a).

**Figure 5:**
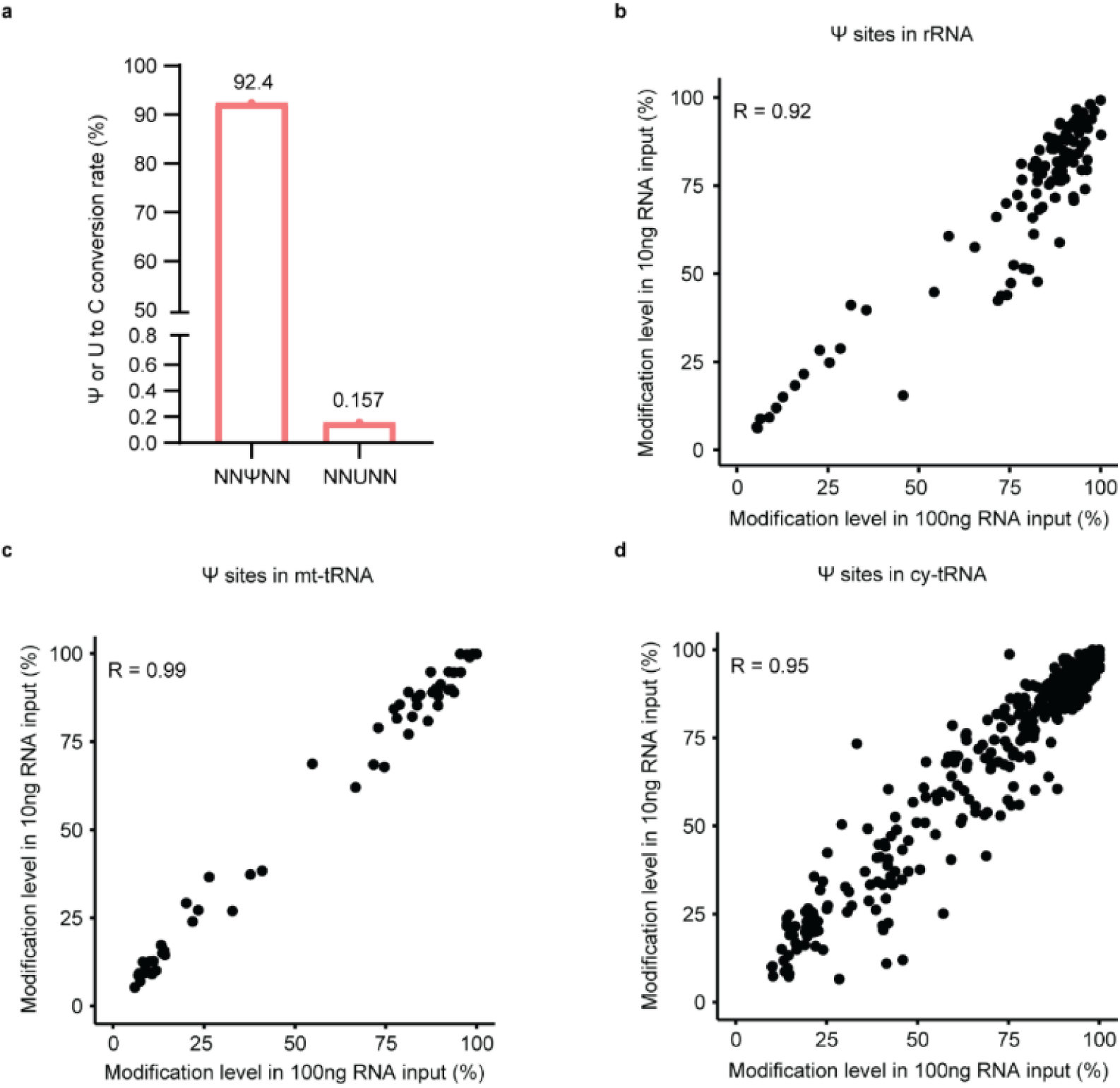
mBACS enables robust and reproducible Ψ quantification from low RNA input. a, Conversion rates of the 30-mer NNΨNN and NNUNN spike-in RNAs in 10ng total RNA input sample detected by mBACS. b, Scatter plot showing correlation of the Ψ modification levels at all identified sites in HCT116 cy-rRNAs detected by mBACS between 100ng RNA input and 10ng RNA input. c, Scatter plot showing correlation of the Ψ modification levels at all identified sites in HCT116 mt-tRNAs detected by mBACS between 100ng RNA input and 10ng RNA input. d, Scatter plot showing correlation of the Ψ modification levels at all identified sites in HCT116 cy-tRNAs detected by mBACS between 100ng RNA input and 10ng RNA input.

Transcriptome-wide Ψ profiles generated from 10 ng of total RNA showed a high correlation with those obtained from 100 ng input (R = 0.92 - 0.99), with highly concordant modification levels across identified Ψ sites, including rRNA, cy-tRNA, and mt-tRNA (Figure 5b-d). Moreover, the majority of Ψ sites identified using the standard input were also detected in the 10 ng samples, indicating that reducing RNA input had minimal impact on detection sensitivity (Figure 5b-d).

Collectively, these results demonstrate that mBACS retains high sensitivity, accuracy, and quantitative reliability even with substantially reduced RNA input amounts. This low-input compatibility further highlights the robustness of mBACS and expands its applicability to precious biological samples and other material-limited experimental settings.

## Discussion

Accurate Ψ profiling has remained challenging because existing chemical and sequencing strategies suffer from incomplete conversion, sequence-context bias, and limited quantitative accuracy^9–14,16,17^. Here, we developed mBACS, a second-generation BACS workflow based on MBA chemistry that substantially improved Ψ conversion efficiency and while maintaining low false-positive rates. The optimized library workflow also reduced the total processing time of mBACS to 2 days compared with 4 days for the original BACS protocol. Improved chemical conversion expanded the detectable pseudouridylation landscape, enabling the identification of previously inaccessible low-stoichiometry Ψ sites, including Ψ65 in mt-tRNAs and Ψ44 in cy-tRNAs. Application of mBACS with 5-FU-treated cells further revealed widespread but highly site-specific pseudouridylation remodeling, demonstrating differential susceptibility of individual Ψ sites and PUS enzymes to pharmacological inhibition. Finally, mBACS enabled accurate and quantitative Ψ profiling from as little as 10 ng of total RNA—approximately the amount recoverable from ∼500 cells— substantially reducing input requirements and broadening its applicability to precious and material-limited samples. Overall, mBACS provides a sensitive, quantitative, robust, and low-input chemical sequencing platform for transcriptome-wide Ψ profiling and expands opportunities to investigate RNA pseudouridylation across diverse biological and therapeutic intervention.

## Experimental methods

### Preparation of synthetic RNAs

A 10-mer Ψ-containing RNA oligonucleotide and synthetic 30-mer RNA spike-ins containing either a single Ψ (NNΨNN) or uridine (NNUNN) were purchased from Integrated DNA Technologies (IDT).

### Reaction of synthetic RNA oligonucleotide with MBA, DPVST, and VTT

The reaction of 10-mer Ψ-containing RNA oligonucleotide with compounds were incubated at 85 °C for 30 min. Following incubation, reaction products were purified using Bio-Spin P-6 desalting columns (Bio-Rad, 7326222) according to the manufacturer’s instructions.

### MALDI-TOF mass spectrometry analysis

Purified reaction products were mixed with a matrix consisting of 2′,4′,6′-trihydroxyacetophenone (THAP) and diammonium citrate and analysed by MALDI-TOF mass spectrometry using a SHIMADZU MALDI instrument.

### Cell Culture

HCT116 human colorectal carcinoma cells (ATCC, CCL-247) were cultured in McCoy’s 5A Modified Medium (Gibco) supplemented with 10% (v/v) fetal bovine serum (FBS; Gibco) and 1% penicillin– streptomycin (Gibco) at 37 °C in a humidified incubator with 5% CO₂. Cell line identity was authenticated by ATCC using short tandem repeat (STR) profiling. All cell lines have been periodically tested in-house for mycoplasma contamination.

### Generation of CRISPR knockout cell lines

Monoclonal TRUB1/PUS10 double-knockout HCT116 cell lines were generated using CRISPR–Cas9- mediated genome editing. Single-guide RNA (sgRNA) sequences targeting TRUB1 and PUS10 were cloned into the PX459 vector. HCT116 cells were transfected using Lipofectamine 3000 Transfection Reagent (Invitrogen) according to the manufacturer’s instructions. Seventy-two hours after transfection, cells were selected with 2 μg mL⁻¹ puromycin (Thermo Fisher Scientific). Monoclonal cell lines were established by limiting dilution, expanded, and validated by Sanger sequencing.

### 5-FU treatment

HCT116 cells were seeded in 12-well plates at a density of 1 × 10⁵ cells per well in biological triplicate and treated with 5-fluorouracil (5-FU) or an equivalent volume of DMSO as a vehicle control for 24, 48, or 72 h at 37 °C in a humidified incubator with 5% CO₂. Cell proliferation was assessed at each time point to evaluate the inhibitory effects of 5-FU. Cells were harvested following 72h of treatment for RNA extraction and subsequent mBACS analysis.

### RNA isolation

Total RNA was extracted using the Quick-RNA Miniprep Kit (Zymo Research, R1054) with on-column DNase I treatment according to the manufacturer’s instructions.

### mBACS library preparation

For standard-input experiments, 100 ng of total RNA supplemented with 0.1% synthetic RNA spike-ins was subjected to mBACS treatment. The reaction mixture of total RNA with compounds in 625 mM phosphate buffer (pH 8.5) was incubated at 85 °C for 30 min. Following incubation, RNA was purified using a Micro Bio-Spin P-6 column (Bio-Rad) followed by a Zymo IC column with RNA Binding Buffer and eluted in 8 μL of nuclease-free water. Untreated control (2 μL) of initial 12 μL of total RNA sample were processed in parallel starting with end-repair step.

RNA samples were subsequently subjected to 3′ and 5′-end repair, then treated RNA was purified using a Zymo IC column and eluted in 9 μL of nuclease-free water.

Sequencing libraries were prepared using the NEBNext® Low-bias Small RNA Library Prep Kit (NEB, R3420S) according to the manufacturer’s instructions. Following reverse transcription and cDNA purification, libraries were amplified by PCR for 11 cycles and purified using the bead cleanup procedure recommended for PCR products without size selection. Final libraries were sequenced on an Illumina NextSeq 1000/2000 platform.

### Low-input mBACS library preparation

For low-input experiments, the mBACS workflow was performed as described above except that 10 ng of total RNA, supplemented with 0.1% synthetic RNA spike-ins, was used as the input material instead of 100 ng. Subsequent chemical treatment, RNA purification, 5′-end repair, library preparation, and sequencing steps were performed using the same conditions as described for the standard-input protocol. Libraries were amplified by PCR for 15 cycles.

### Data preprocessing

Raw sequencing reads were processed using Cutadapt (v4.9)^31^ to remove adaptor sequences, low-quality bases (quality score <20), and reads shorter than 18 nt. Paired-end reads were subsequently merged using fastp (v0.23.2)^32^.

### Read alignment

Cleaned reads were first mapped to synthetic spike-ins and rRNA references using bowtie2 (v.2.4.4)^33^. The unaligned reads were subsequently mapped to tRNA references. High-confidence human tRNA sequences (hg38) were downloaded from GtRNAdb^34^. Only non-redundant tRNA sequences were kept and appended with a “3’-CCA” end.

The aligned reads were then filtered and sorted using SAMtools (v.1.16.1)^35^. For synthetic spike-ins and rRNAs, only reads with MAPQ ≥ 10 were kept. For tRNAs, only reads with MAPQ ≥ 1 were kept. Reads with correct strand information were kept. Finally, mutations were counted by SAMtools mpileup (v.1.16.1)^35^ and cpup (v.0.1.0) (https://github.com/y9c/cpup).

### Calling Ψ sites

Ψ modification levels were quantified from T-to-C conversion events. Raw conversion rates were calculated as the fraction of C reads among all T and C reads (C/(T + C)). Modification levels were then estimated using a linear correction model: Ψ modification level = (R − F)/(C − F), where R is the observed conversion rate, F is the sequence motif-specific background conversion rate determined from the corresponding negative control and C is the motif-specific conversion efficiency measured using NNΨNN spike-ins.

High-confidence Ψ sites in non-coding RNAs were required to satisfy the following criteria: a minimum sequencing depth of 20 reads in both treated and untreated libraries; a background conversion rate ≤0.01 or no more than two T-to-C mismatches in the untreated library; a background T-to-R (A or G) substitution frequency ≤0.10; and an estimated Ψ modification level of at least 5%. For calling cy-tRNA Ψ sites, the above criteria were modified to require a Ψ modification level ≥ 10% in at least one of two replicates. A P value was calculated for each site using the motif-specific false-positive rates and then adjusted following the Benjamini–Hochberg procedure; the adjusted P value is required to be <0.001.

### Statistics and reproducibility

Statistical analyses were performed using R (v4.0.3). Unless otherwise specified, experiments included two or three independent biological replicates. Differences between two groups were evaluated using paired, two-sided *t*-tests. Data were assumed to follow a normal distribution, although this assumption was not formally assessed. Individual observations are displayed in the corresponding box plots.

Sample sizes were not determined using statistical methods but were comparable to those used in our previous studies. No samples or data points were excluded from the analyses. Experiments were not randomized, and investigators were not blinded during data collection or analysis because all samples were processed using the same standardized experimental workflow and conditions.

### Published data

Related published data were downloaded from the Gene Expression Omnibus (GEO) database: TRUB1- KO, PUS7-KO and PUS1-KO HeLa cells (GSE241849)^18,19^ and PUS10-KO HCT116 cells (GSE285932).

## Acknowledgements

The authors would like to thank Mengjie Li from Dr. Medhipour’s lab (Ludwig Institute for Cancer Research) for sharing HCT116 cells, TRUB1 sgRNAs and primers. This work was funded by the Ludwig Institute for Cancer Research (to C.-X.S.). C.-X.S. lab is also supported by the National Institute for Health Research (NIHR) and Oxford Biomedical Research Centre (BRC). L.K. are supported by China Scholarship Council (CSC). T.G. and P.E.B. are supported by the Alzheimer’s Research UK (grant no. ARUK-2021DDI-OX). Computation used the Oxford Biomedical Research Computing (BMRC) facility, a joint development between the Wellcome Centre for Human Genetics and the Big Data Institute supported by Health Data Research UK and the NIHR Oxford Biomedical Research Centre. The views expressed are those of the author(s) and not necessarily those of the NHS, the NIHR, or the Department of Health.

## Author contributions

C.-X.S., F.F., and G.P. conceived and designed the study. F.F. developed and optimised the mBACS chemistry. G.P. developed and optimised the library preparation workflow for mBACS. F.F. and G.P. jointly generated experimental datasets. L.K. genereted HCT116 DKO cells and performed the computational analyses, with input from C.-X.S., F.F., and G.P.. T.G., from Prof. Brennan’s laboratory, conducted the LC-MS, HPLC, and NMR experiments. C.-X.S., F.F., G.P., and L.K. interpreted the results and contributed to writing and revising the manuscript.

## Competing interests

All authors declare no competing interests.

## Notes

### Competing Interest Statement

The authors have declared no competing interest.

