## supplemental notes and figure for "mBACS enhances pseudouridine profiling through efficient and robust chemical conversion"

### Supplementary Note 1: Reaction of MBA with pseudouridine

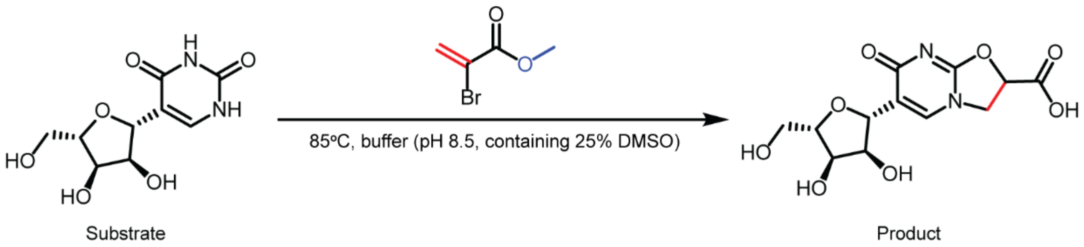


For structural characterization of the MBA reaction product: methyl α-bromoacrylate (13.2 mg, 0.0800 mmol; Sigma-Aldrich, 4519-46-4) dissolved in 500 μL DMSO was added to pseudouridine (10 mg, 0.0410 mmol; Fluorochem, 1445-07-4) dissolved in 1.5 mL of 0.833 M phosphate buffer. The reaction mixture was incubated at 85 °C for 30 min, after which DMSO was removed under reduced pressure. Excess phosphate salts were removed by trituration with MeOH, and the filtrate was concentrated *in vacuo*. The crude product was purified by preparative HPLC (0 to 15% MeCN in water modified with 0.1% formic acid) to afford the product as a colourless oil (6.58 mg, 51.1% yield).

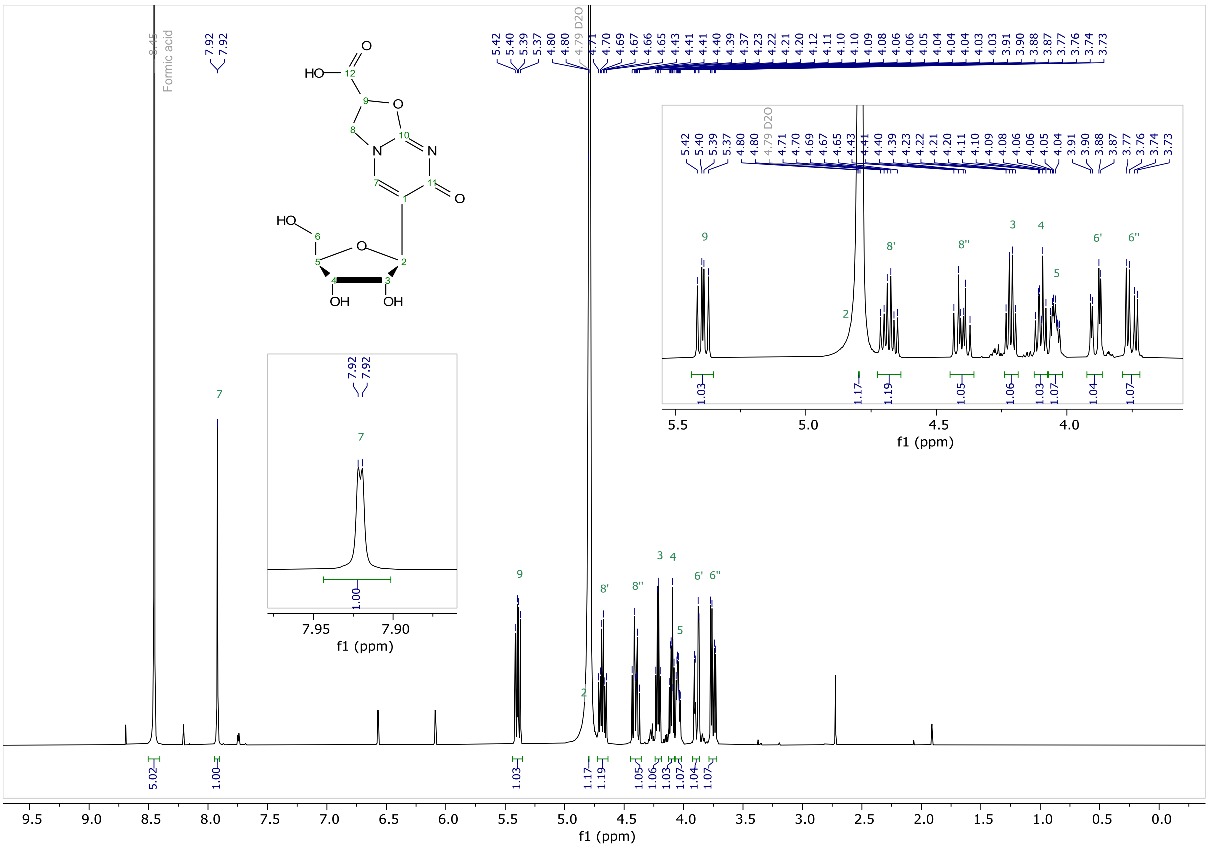


[1]H NMR (400 MHz, $D_2O$) δ 7.92 (d, *J* = 1.1 Hz, 1H, H7), 5.39 (dd, *J* = 10.2, 7.1 Hz, 1H, H9), 4.80 – 4.79 (m, 1H, H2), 4.68 (td, *J* = 10.3, 5.5 Hz, 1H, H8’), 4.40 (dt, *J* = 10.4, 7.1 Hz, 1H, H8’’), 4.21 (q, *J* = 4.9 Hz, 1H, H3), 4.10 (dt, *J* = 6.5, 4.9 Hz, 1H, H4), 4.05 (ddd, *J* = 6.6, 4.6, 2.9 Hz, 1H, H5), 3.89 (dd, *J* = 12.6, 2.8 Hz, 1H, H6’), 3.75 (dd, *J* = 12.6, 4.7 Hz, 1H, H6’’).

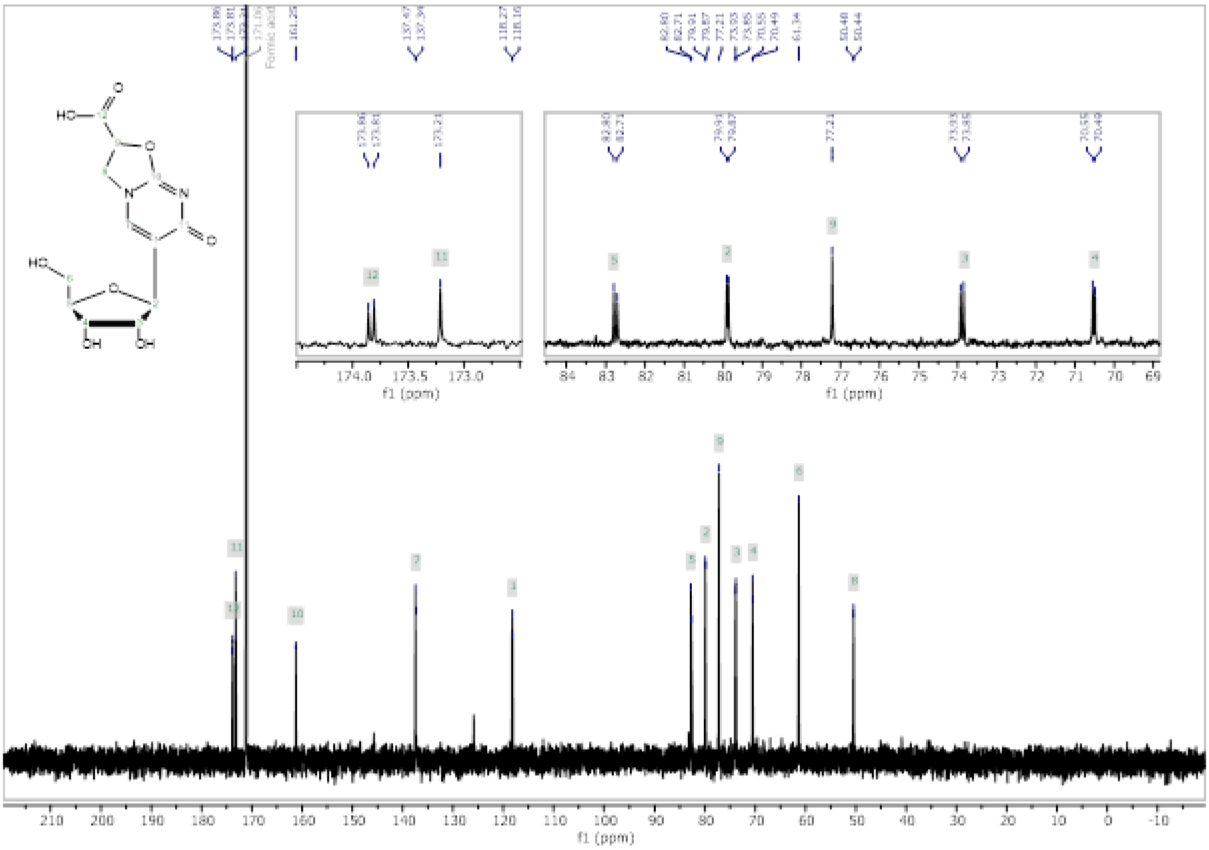

$^{13}C$ NMR (101 MHz, $D_2O$) δ 173.9 (C12), 173.8 (C11a), 173.2 (C11b), 161.2 (C10), 137.5 (C7a), 137.3 (C7b), 118.3 (C1a), 118.2 (C1b), 82.8 (C5a), 82.7 (C5b), 79.9 (C2a), 79.9 (C2b), 77.2 (C9), 73.9 (C3a), 73.8 (C3b), 70.5 (C4a), 70.5 (C4b), 61.3 (C6), 50.5 (C8a), 50.4 (C8b). Diastereomer peaks are arbitrarily assigned as “a” or “b”.

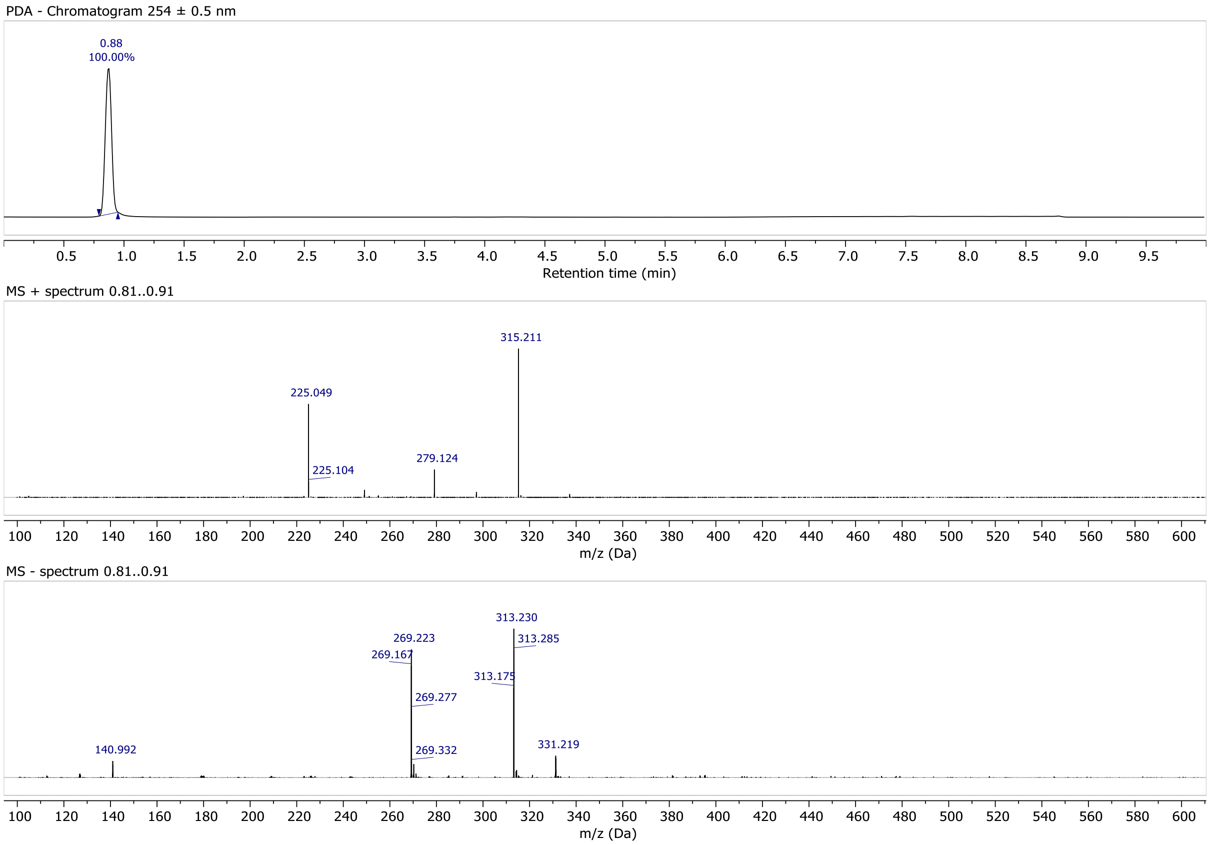


MS (ESI) for $[M + H]^+$ ($C_{12}H_{15}N_2O_8^+$): calcd m/z 315.082; found m/z 315.211. MS (ESI) for $[M - H]^-$ ($C_{12}H_{13}N_2O_8^-$): calcd m/z 313.068; found m/z 313.230. LC–MS: Rt = 0.88, 100% purity at 254 nm.

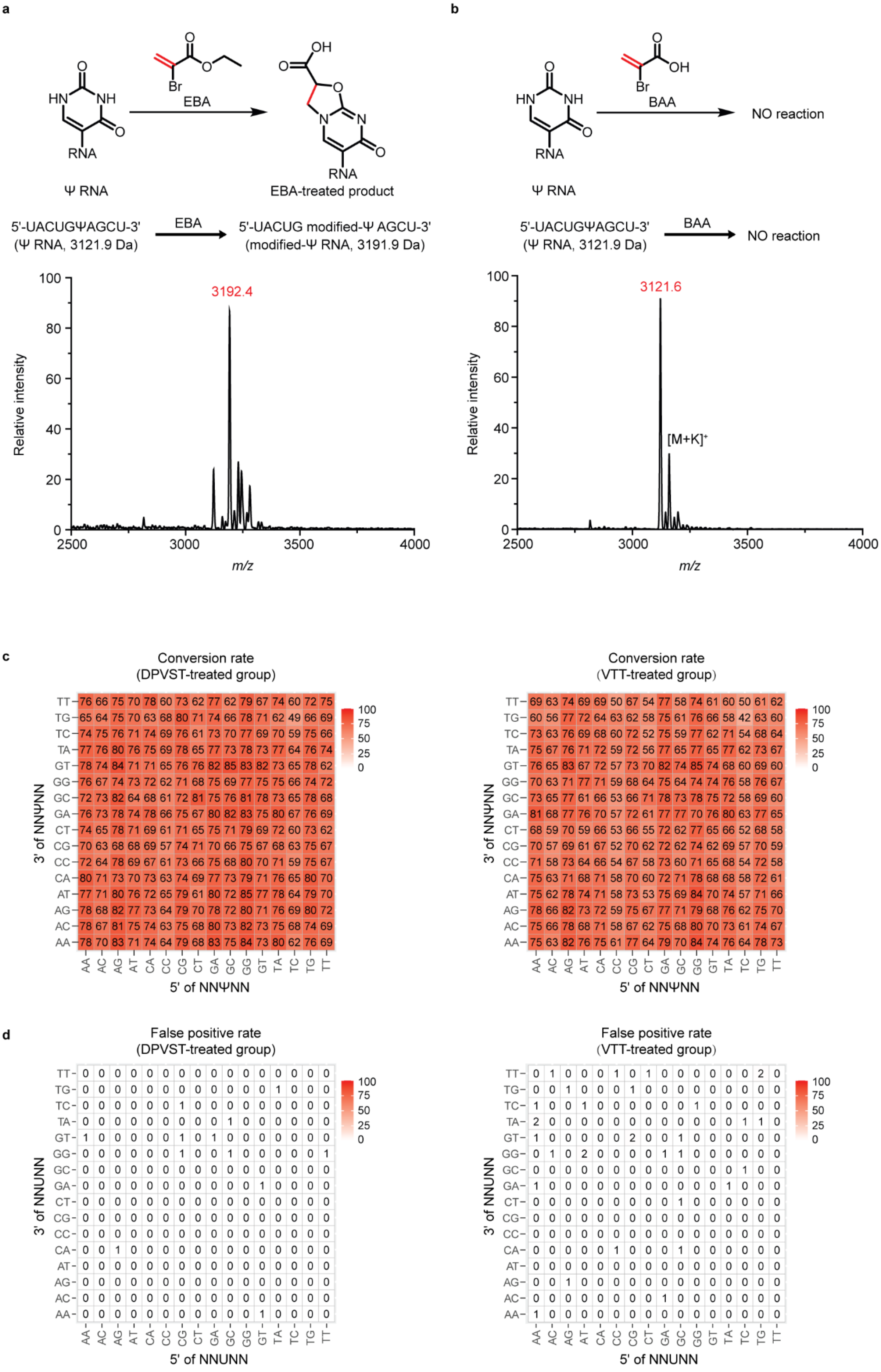
a
EBA
Br
OH
O
HN
NH
N
RNA
Ψ RNA
EBA-treated product
5'-UACUGΨAGCU-3'
(Ψ RNA, 3121.9 Da)
EBA
5'-UACUG modified-Ψ AGCU-3'
(modified-Ψ RNA, 3191.9 Da)
3192.4
Relative intensity
m/z
100
80
60
40
20
0
2500
3000
3500
4000
b
BAA
NO reaction
5'-UACUGΨAGCU-3'
(Ψ RNA, 3121.9 Da)
BAA
NO reaction
3121.6
[M+K]+
c
Conversion rate
(DPVST-treated group)
TT 76 66 75 70 78 60 73 62 77 62 79 67 74 60 72 75
TG 65 64 75 70 63 68 80 71 74 66 78 71 62 49 66 69
TC 74 75 76 71 74 69 76 61 73 70 77 69 70 59 75 66
TA 77 76 80 76 75 69 78 65 77 73 78 73 77 64 76 74
GT 78 74 84 71 71 65 76 76 82 85 83 82 73 65 78 62
GG 76 67 74 73 72 62 71 68 75 69 77 75 75 66 74 72
GC 72 73 82 64 68 61 72 81 75 76 81 78 73 65 78 68
GA 76 73 78 74 78 66 75 67 80 82 83 75 80 67 76 69
CT 74 65 78 71 69 61 71 65 75 71 79 69 72 60 73 62
CG 70 63 68 68 69 57 74 71 70 66 75 67 68 63 75 67
CC 72 64 78 69 67 61 73 66 75 68 80 70 71 59 75 67
CA 80 71 73 70 73 63 74 69 77 73 79 71 76 65 80 70
AT 77 71 80 76 72 65 79 61 80 72 85 77 78 64 79 70
AG 78 68 82 77 73 64 79 70 78 72 80 71 76 69 80 72
AC 78 67 81 75 74 63 75 68 80 73 82 73 75 68 74 69
AA 78 70 83 71 74 64 79 68 83 75 84 73 80 62 76 69
AA AC AG AT CA CC CG CT GA GC GG GT TA TC TG TT
3' of NNΨNN
5' of NNΨNN
100
75
50
25
0
Conversion rate
(VTT-treated group)
TT 69 63 74 69 69 50 67 54 77 58 74 61 60 50 61 62
TG 60 56 77 72 64 63 62 58 75 61 76 66 58 42 63 60
TC 73 63 76 69 68 60 72 52 75 59 77 62 71 54 68 64
TA 75 67 76 71 72 59 72 56 77 65 77 65 77 62 73 67
GT 76 65 83 67 72 57 73 70 82 74 85 74 68 60 69 60
GG 70 63 71 77 71 59 68 64 75 64 74 74 76 58 76 67
GC 73 65 77 61 66 53 66 71 78 73 78 75 72 58 69 60
GA 81 68 77 76 70 57 72 61 77 77 70 76 80 63 77 65
CT 68 59 70 59 66 53 66 55 72 62 77 65 66 52 68 58
CG 70 57 69 61 67 52 70 62 72 62 74 62 69 57 70 59
CC 71 58 73 64 66 54 67 58 73 60 71 65 68 54 72 58
CA 75 63 71 68 71 58 70 60 72 71 78 68 68 58 72 61
AT 75 62 78 74 71 58 73 53 75 69 84 70 74 57 71 66
AG 78 66 82 73 72 59 75 67 77 71 79 68 76 62 75 70
AC 75 66 78 68 71 58 67 62 75 68 80 70 73 61 74 67
AA 75 63 82 76 75 61 77 64 79 70 84 74 76 64 78 73
AA AC AG AT CA CC CG CT GA GC GG GT TA TC TG TT
3' of NNΨNN
5' of NNΨNN
d
False positive rate
(DPVST-treated group)
TT 0 0 0 0 0 0 0 0 0 0 0 0 0 0 0 0
TG 0 0 0 0 0 0 0 0 0 0 0 0 1 0 0 0
TC 0 0 0 0 0 0 1 0 0 0 0 0 0 0 0 0
TA 0 0 0 0 0 0 0 0 0 1 0 0 0 0 0 0
GT 1 0 0 0 0 0 1 0 1 0 0 0 0 0 0 0
GG 0 0 0 0 0 0 1 0 0 1 0 0 0 0 0 1
GC 0 0 0 0 0 0 0 0 0 0 0 0 0 0 0 0
GA 0 0 0 0 0 0 0 0 0 0 0 1 0 0 0 0
CT 0 0 0 0 0 0 0 0 0 0 0 0 0 0 0 0
CG 0 0 0 0 0 0 0 0 0 0 0 0 0 0 0 0
CC 0 0 0 0 0 0 0 0 0 0 0 0 0 0 0 0
CA 0 0 1 0 0 0 0 0 0 0 0 0 0 0 0 0
AT 0 0 0 0 0 0 0 0 0 0 0 0 0 0 0 0
AG 0 0 0 0 0 0 0 0 0 0 0 0 0 0 0 0
AC 0 0 0 0 0 0 0 0 0 0 0 0 0 0 0 0
AA 0 0 0 0 0 0 0 0 0 0 0 1 0 0 0 0
AA AC AG AT CA CC CG CT GA GC GG GT TA TC TG TT
3' of NNUNN
5' of NNUNN
False positive rate
(VTT-treated group)
TT 0 1 0 0 0 1 0 1 0 0 0 0 0 0 2 0
TG 0 0 1 0 0 0 1 0 0 0 0 0 0 0 0 0
TC 1 0 0 1 0 0 0 0 0 0 1 0 0 0 0 0
TA 2 0 0 0 0 0 0 0 0 0 0 0 0 1 1 0
GT 1 0 0 0 0 0 2 0 0 1 0 0 0 0 0 0
GG 0 1 0 2 0 0 0 0 1 1 0 0 0 0 0 0
GC 0 0 0 0 0 0 0 0 0 0 0 0 0 1 0 0
GA 1 0 0 0 0 0 0 0 0 0 0 0 1 0 0 0
CT 0 0 0 0 0 0 0 0 0 1 0 0 0 0 0 0
CG 0 0 0 0 0 0 0 0 0 0 0 0 0 0 0 0
CC 0 0 0 0 0 0 0 0 0 0 0 0 0 0 0 0
CA 0 0 0 0 0 1 0 0 0 1 0 0 0 0 0 0
AT 0 0 0 0 0 0 0 0 0 0 0 0 0 0 0 0
AG 0 0 1 0 0 0 0 0 0 0 0 0 0 0 0 0
AC 0 0 0 0 0 0 0 0 1 0 0 0 0 0 0 0
AA 1 0 0 0 0 0 0 0 0 0 0 0 0 0 0 0
AA AC AG AT CA CC CG CT GA GC GG GT TA TC TG TT
3' of NNUNN
5' of NNUNN

**Supplementary Figure 1: Chemical characterization of EBA and BAA and additional performance evaluation of DPVST and VTT.**

a, Reaction scheme (top) and MALDI-TOF mass spectra (bottom) of the reaction between EBA and a 10-mer Ψ-containing RNA oligonucleotide. b, Reaction scheme (top) and MALDI-TOF mass spectra (bottom) of the reaction between BAA and a 10-mer Ψ-containing RNA oligonucleotide. Calculated masses indicated in black, and detected masses indicated in red. c, Conversion rates of the 30-mer NNΨNN spike-in RNA obtained following treatment with DPVST (left) and VTT (right). d, False positive rates of the 30-mer NNUNN spike-in RNA following treatment with DPVST (left) and VTT (right).

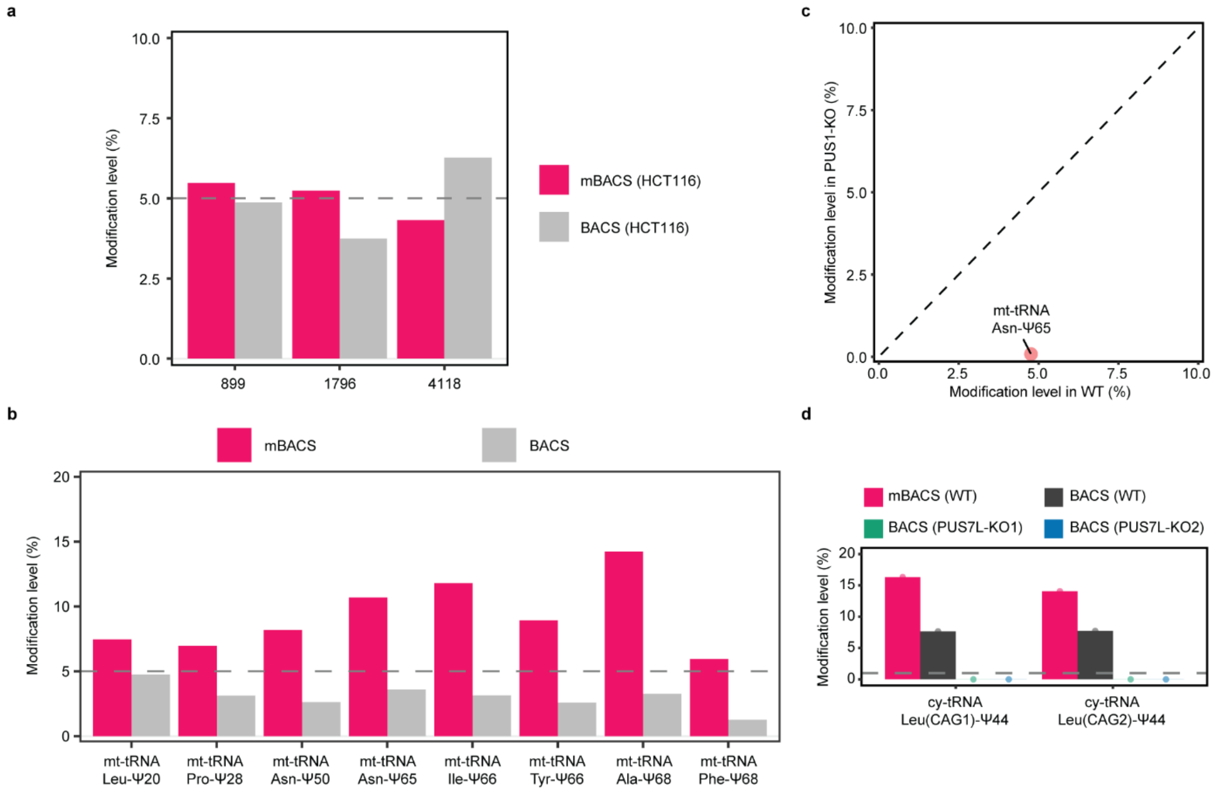


**Supplementary Figure 2: Extended validation of newly identified pseudouridine sites detected by mBACS.**

a, Modification levels of Ψ899, Ψ1795, and Ψ4118 sites in HCT116 cy-rRNAs between mBACS (red) and BACS (grey). b, Modification levels in eight mBACS-unique sites in mt-tRNAs between mBACS (red) and previously reported BACS (grey), including Leu-Ψ20, Pro-Ψ28, Asn-Ψ50, Asn-Ψ65, Ile-Ψ66, Tyr-Ψ66, Ala-Ψ68, and Phe-Ψ68. c, Scatter plot showing modification levels of Ψ65 in mt-tRNAs in HeLa WT and HeLa PUS1-KO. d, Modification levels of two Ψ44 sites in cy-tRNAs between HCT116 WT (red as mBACS and black as BACS), HCT116 PUS7L-KO1 (green), and HCT116 PUS7L-KO2 (blue). BACS data refer to previous dataset[1,2].

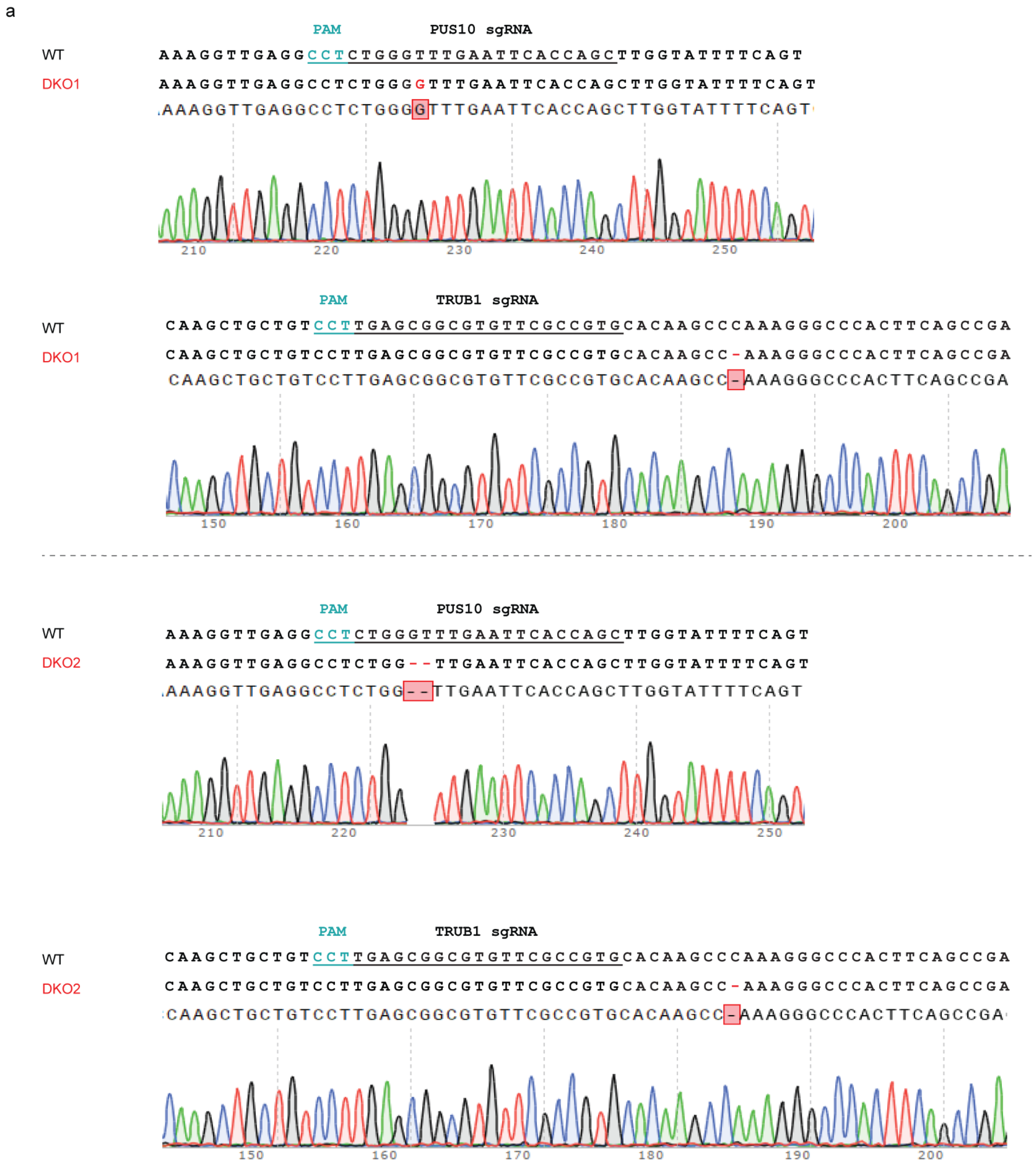


**Supplementary Figure 3: Validation of HCT116 DKO cells.**

a, Sanger sequencing of the two DKO cell clones, showing CRISPR-induced indels in HCT116 cells.

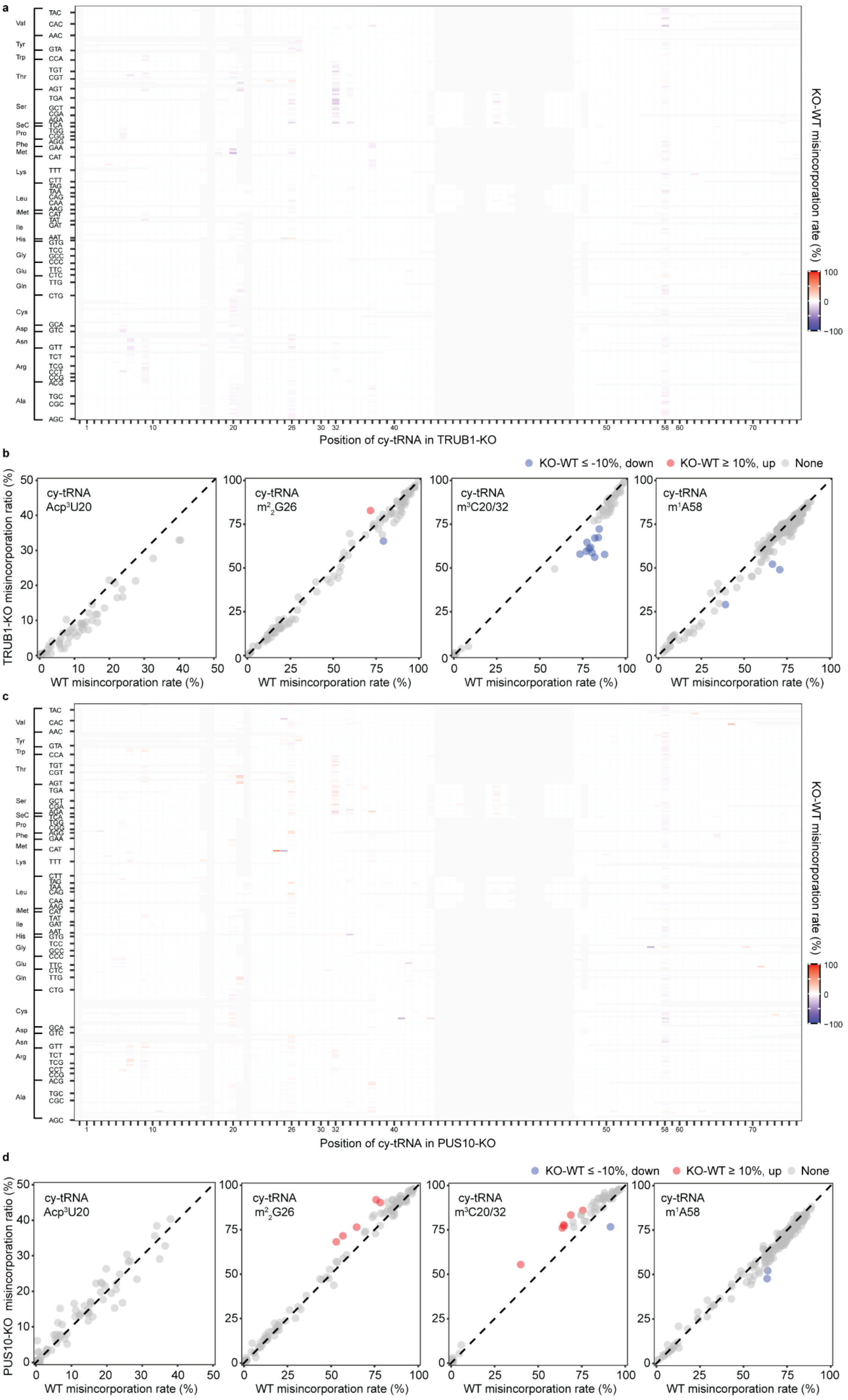
a
KO-WT misincorporation rate (%)
Position of cy-tRNA in TRUB1-KO
b
KO-WT ≤ -10%, down
KO-WT ≥ 10%, up
None
TRUB1-KO misincorporation ratio (%)
cy-tRNA Acp³U20
cy-tRNA m²₂G26
cy-tRNA m³C20/32
cy-tRNA m¹A58
WT misincorporation rate (%)
c
KO-WT misincorporation rate (%)
Position of cy-tRNA in PUS10-KO
d
KO-WT ≤ -10%, down
KO-WT ≥ 10%, up
None
PUS10-KO misincorporation ratio (%)
cy-tRNA Acp³U20
cy-tRNA m²₂G26
cy-tRNA m³C20/32
cy-tRNA m¹A58
WT misincorporation rate (%)

**Supplementary Figure 4: Neither TRUB1 nor PUS10 KO alone shows dramatic changes in other modifications in cy-tRNAs.**

a, Heatmap showing the differential misincorporation ratio of tRNA bases upon TRUB1 KO. The x axis represents the canonical position of each aligned cy-tRNAs isodecoders. The y axis represents different cy-tRNA isodecoders. b, Scatter plots illustrating the misincorporation rate in $acp^3U20$, $m^2{}_2G26$, $m^3C20/32$, $m^1A58$ in HeLa WT and HeLa TRUB1 KO. c, Heatmap showing the differential misincorporation ratio of tRNA bases upon PUS10 KO. The x axis represents the canonical position of each aligned cy-tRNAs isodecoders. The y axis represents different cy-tRNA isodecoders. d, Scatter plots illustrating the misincorporation rate of $acp^3U20$, $m^2{}_2G26$, $m^3C20/32$, $m^1A58$ in HCT116 WT and HCT116 PUS10 KO**.**

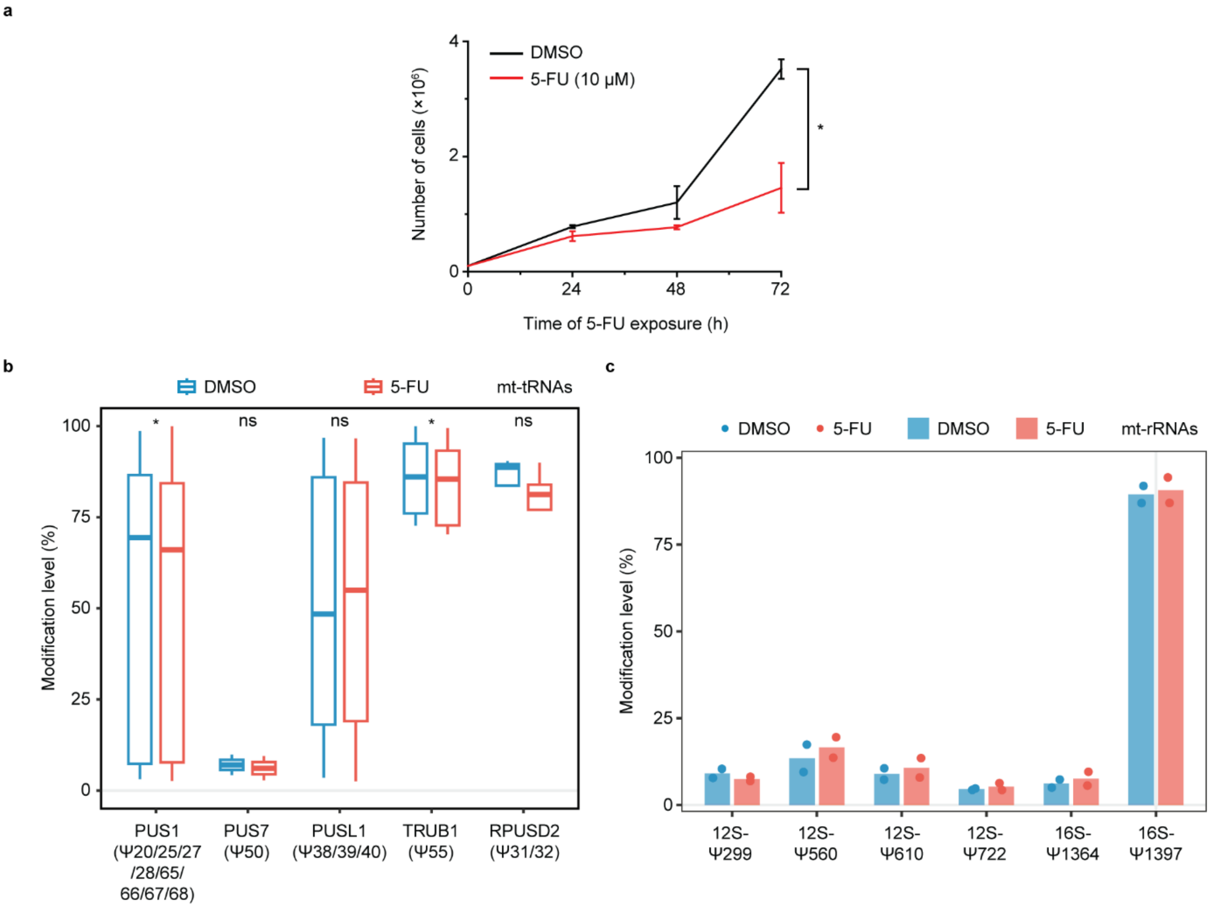


**Supplementary Figure 5: 5-FU inhibits cell growth and alters mitochondrial tRNA pseudouridylation**

a, Time-dependent growth curves of HCT116 treated with DMSO (control) or 5-FU (10 μM). b, Modification levels at identified sites in HCT116 mt-tRNAs between DMSO and 5-FU treated group after 72h of treatment. Boxes represent the percentiles with a line at the median (PUS1: n=28 Ψ sites; PUS7: n=2 Ψ sites; PUSL1: n=16 Ψ sites; TRUB1: n=7 Ψ sites; RPUSD2: n=4 Ψ sites). c, Modification levels of identified Ψ sites in mt-rRNAs between DMSO and 5-FU treated group after 72h of treatment. *P* values were calculated using paired, two-tailed t-test. ns, $P >= 0.05$; * $P < 0.05$; ** $P < 0.01$; *** $P < 0.001$ and **** $P < 0.0001$.
